# Mechanical convergence in mixed populations of mammalian epithelial cells

**DOI:** 10.1101/2024.02.18.580863

**Authors:** Estelle Gauquelin, Keisuke Kuromiya, Toshinori Namba, Keisuke Ikawa, Yasuyuki Fujita, Shuji Ishihara, Kaoru Sugimura

## Abstract

Tissues consist of cells with different molecular and/or mechanical properties. Measuring the forces and stresses in mixed-cell populations is essential for understanding the mechanisms by which tissue development, homeostasis, and disease emerge from the cooperation of distinct cell types. However, many previous studies have primarily focused their mechanical measurements on dissociated cells or aggregates of a single-cell type, leaving the mechanics of mixed-cell populations largely unexplored. In the present study, we aimed to elucidate the influence of interactions between different cell types on cell mechanics by conducting *in situ* mechanical measurements on a monolayer of mammalian epithelial cells. Our findings revealed that while individual cell types displayed varying magnitudes of traction and intercellular stress before mixing, these mechanical values shifted in the mixed monolayer, becoming nearly indistinguishable between the cell types. Moreover, by analyzing a mixed-phase model of active tissues, we identified physical conditions under which such mechanical convergence is induced. Overall, the present study underscores the importance of *in situ* mechanical measurements in mixed-cell populations to deepen our understanding of the mechanics of multi-cellular systems.

## 1 Introduction

Tissues in health and disease are composed of cells with different molecular and/or mechanical characteristics. The development, homeostasis, and dysfunction of tissues arise from cell-cell communications within such heterotypic tissue environments [1–7].

Distinct cell types within a tissue communicate through mechanical forces as well as biochemical signaling [2, 4, 8–13]. For instance, stress transmission from one cell type to another can induce the remodeling of the extracellular matrix [14]. The precise formation of compartment boundaries relies on the differences in cell adhesion strength [15, 16]. Forces between oncogenic cells and their neighboring normal cells determine which will be eliminated from the tissue, a process driven by short-range cell-cell communications known as cell competition [17–19]. Identifying the mechanisms through which mechanical communication between diverse cell types controls tissue dynamics will deepen our understanding of the physics of multicellular systems.

To address this, it is crucial to measure the forces and stresses in mixed-cell populations. In many previous studies, mechanical measurements have focused on dissociated cells or aggregates of a single-cell type [20–24]. This can be attributed, in part, to the challenges associated with *in situ* measurements of intercellular stress, which were difficult to achieve until recent advancements [25, 26]. Often, results from single-cell-type measurements have been used to interpret the dynamics of mixed-cell populations under the assumption that cell mechanics remain consistent regardless of the presence of other cell types. However, this assumption has not been rigorously tested, leaving the mechanics of mixed-cell populations largely unexplored.

Bayesian inversion stress microscopy (BISM) offers a mean to measure intercellular stress within a monolayer of cultured cells [27]. Traction force microscopy (TFM) is used to measure the traction exerted by the cells on a substrate [28]. BISM then computes the stress tensor from the measured traction. This method has been validated using synthetic data and successfully utilized to characterize stress-dependent cellular behaviors such as apoptosis and rigidity sensing [27, 29–32]. Using BISM, we recently quantified intercellular stress under conditions that initiate cell competition between oncogenic cells and their surrounding normal counterparts [33]. Our findings indicate that normal cells decreased the isotropic component of intercellular stress when co-cultured with oncogenic RasV12 cells. This alteration in isotropic intercellular stress subsequently triggers down-stream biochemical signaling, leading to the apical extrusion of RasV12 cells. These findings highlight the significance of conducting mechanical measurements directly within mixed-cell populations to shed light on the mechanical regulation of tissue homeostasis. However, our previous study did not analyze the isotropic stress in RasV12 cells. Moreover, it remains to be determined whether other mechanical quantities, such as deviatoric stress and cell velocity, undergo changes similar to isotropic stress during cell competition.

In the present study, we aimed to elucidate the influence of the interactions between different cell types on epithelial mechanics. Taking the advantage of BISM, we conducted *in situ* mechanical measurements in mixed populations of wild type and genetically modified Madin-Darby canine kidney (MDCK) cells. Our results demonstrated that although individual cell types exhibited distinct levels of traction and intercellular stress before mixing, in the mixed monolayer, these mechanical values were nearly identical between the cell types. This convergence of traction and intercellular stress occurs in the context of both cell competition and differential cell adhesion. Finally, by analyzing a mixed-phase model of active tissues, we showed that the degree of phase separation influences whether two cell types, each with inherently distinct mechanical properties, merge into a single cohesive entity upon mixing.

## 2 Results

To study the cell mechanics in mixed populations, we conducted *in situ* mechanical measurements in a monolayer of cultured epithelial cells. TFM [28] and BISM [27] were employed to quantify the traction exerted by cells on the substrate and the stress tensors within the monolayer (Materials and Methods).

The cells exert tractions 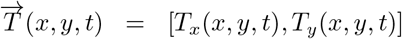 on the substrate. Over each time frame for each monolayer, we calculated the total magnitude of the force 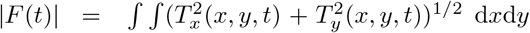. Subsequently, the value |*F* (*t*)| was averaged between experiments performed with the same cell lines in order to obtain traction values plotted over time. Using BISM, we obtained the intercellular stress tensor 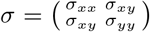 which is symmetrical, through the relationship div 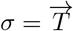.

The isotropic stress *σ*_*iso*_ is a scalar value calculated by taking the trace of the stress tensor *σ* as 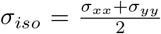. It is the opposite of the pressure inside the tissue, meaning that negative values of *σ*_*iso*_ represent compression in the tissue, and positive values indicate that the tissue is under tension. The deviatoric stress *σ*_*dev*_ is a tensor defined as *σ*_*dev*_ = *σ* − *σ*_*iso*_*I* where *I* is the identity matrix. As *σ*_*dev*_ is a 2-D symmetrical and trace-less matrix, it possesses two eigenvalues, *λ*_*dev*_ and −*λ*_*dev*_, that have the same magnitude but opposite signs. We adopted |*λ*_*dev*_| as the magnitude of the deviatoric stress. Similar to the traction values, the magnitudes of isotropic and deviatoric stresses were averaged first over space and then over experiments.

The experiments were performed using either a single-cell line or a mixture of two cell lines (Fig. 1). By comparing the mechanical behaviors of cells in each type of experiment, we aimed to elucidate how the interactions between different cell types influence cell mechanics.

**Fig. 1.**
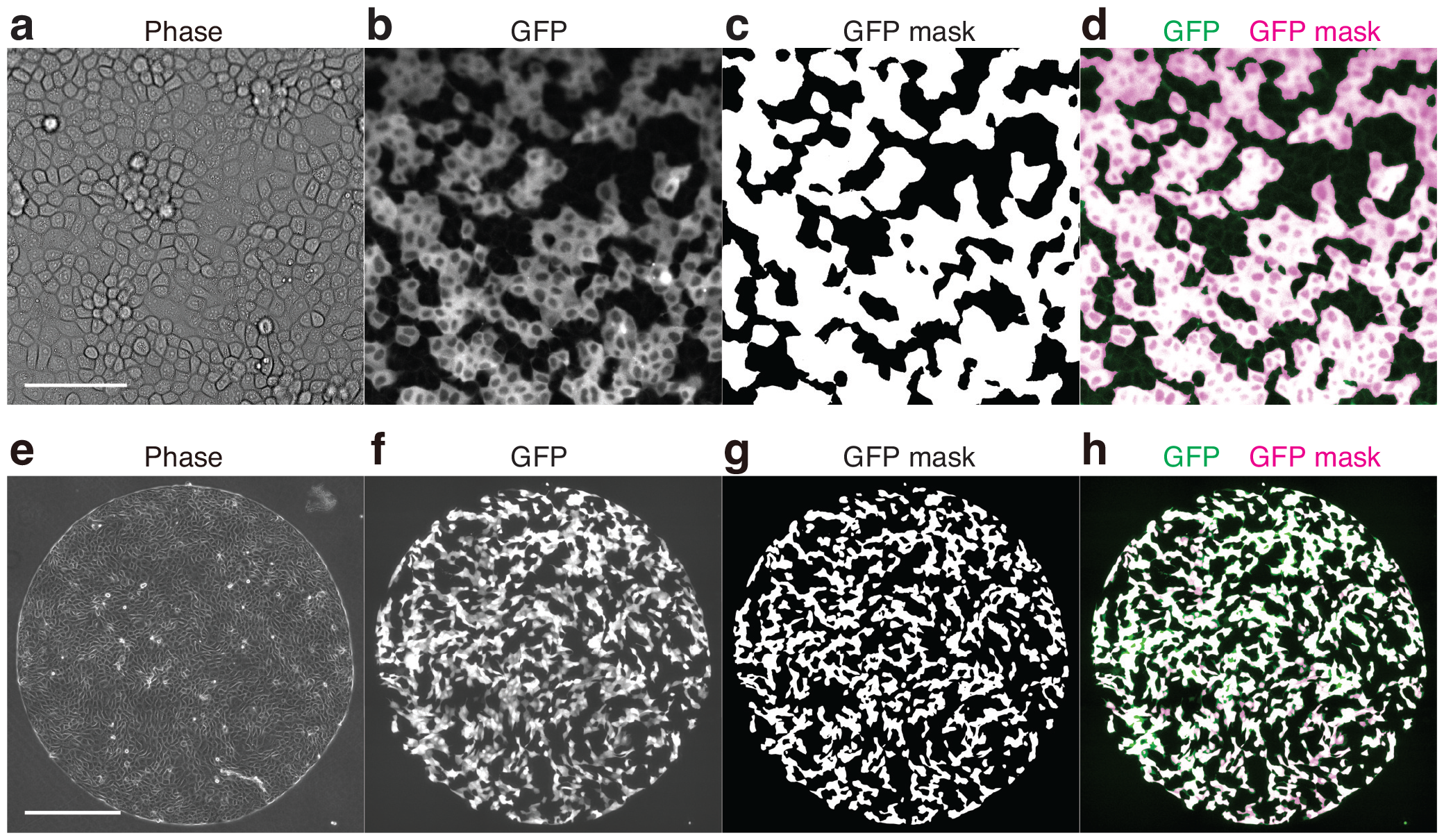
Comparison between GFP fluorescent signal images and their corresponding GFP masks of mixed monolayers. (a–d) Comparison for a WT+RasV12 monolayer comprising GCaMP WT MDCK cells and Myc-tagged RasV12-induced MDCK cells. Phase contrast (a), GFP (GCaMP) (b), GFP mask (c), and the overlay of the GFP image (green) with the GFP mask (magenta) (d) at 6 h after the induction of RasV12 expression. (e-h) Comparison for a WT+E-cad KO monolayer comprising GFP WT MDCK-II cells and non-tagged E-cad KO MDCK-II cells. Phase contrast (e), GFP (f), GFP mask (g), and the overlay of the GFP image (green) with the GFP mask (magenta) (h) at 24 h after cell seeding. Scale bars: 100 μm (a) and 500 μm (e).

### 2.1 WT cells and RasV12 cells respectively decreased and increased traction and intercellular stress in the presence of the other, converging to comparable levels

In multicellular tissues, cells compete for limited resources, such as space and growth factors [34, 35]. Depending on differences in cellular fitness, this cell competition can lead to the elimination of unwanted cells, including those with reduced differentiation potential or those harboring oncogenic mutations [6, 36].

To assess traction and intercellular stress during the early phase of cell competition between normal and oncogenic cells, we mixed wild type (WT) MDCK cells with RasV12 inducible MDCK cells at a 1:1 ratio and seeded them on collagen-coated soft substrates (Fig. 1a–d; Materials and Methods) [33]. We conducted the mechanical measurement at 6-10 hours (h) after the induction of RasV12 expression. Because the extrusion of RasV12 cells starts at considerably later stages [33, 37], both the WT and RasV12 cells remained in a cohesive monolayer during this period. Throughout the duration of our measurement, the number of cells in the monolayers increased only by around 10%. The change in the cell density was thus negligible.

The WT cells exerted higher traction to their substrate and exhibited larger isotropic and deviatoric stresses over time than the RasV12-induced cells when cultured separately (light green and red lines in Fig. 2a, c, e; Video A1). In the monolayer of a mixture of WT and RasV12-induced cells (hereafter referred to as the WT+RasV12 monolayer), the values of traction and isotropic stress fell between those in the WT and RasV12 monolayers (blue lines in Fig. 2a, c; Video A1). The magnitude of deviatoric stress was comparable between the WT and WT+RasV12 monolayers (light green and blue lines in Fig. 2e). We also verified that the variance in mechanical quantities observed across the different types of experiments was not influenced by the size of the region of interest (ROI) (Fig. A1a, c, e, g).

**Fig. 2.**
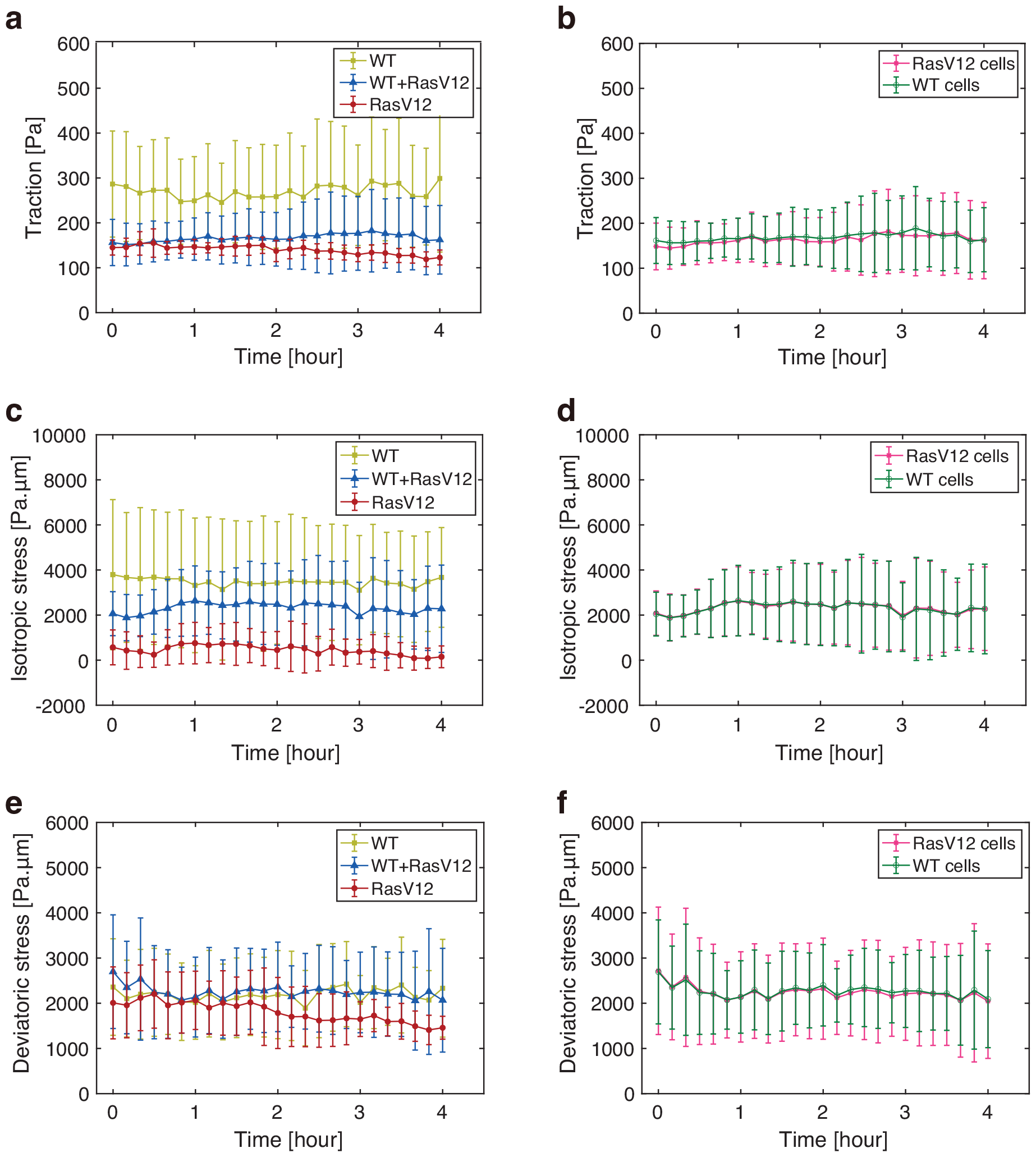
Quantification of traction, isotropic stress, and deviatoric stress during the early phase of cell competition in MDCK cells. (a, b) Magnitude of the traction exerted by the cells in a function of time, depending on the experiment types (a) and on the cell types within the WT+RasV12 co-culture experiments (b) (n = 3 for each plot). 0 hour corresponds to 6 h after the induction of RasV12 expression. In (a), WT represents the co-culture of GCaMP WT and non-tagged WT MDCK cells (light green), WT+RasV12 represents the co-culture of GCaMP WT and Myc-tagged RasV12-induced MDCK cells (blue), and RasV12 represents the co-culture of CMFDA-stained RasV12-induced and unstained RasV12 MDCK cells (red). In (b), the traction magnitude of each cell type (WT: green, RasV12: magenta) in the WT+RasV12 co-culture experiments is plotted. (c, d) Isotropic stress within the monolayers in a function of time, depending on the experiment type (c) and on the cell type within the WT+RasV12 co-culture experiments (d), as shown in (a) and (b). (e, f) Magnitude of deviatoric stress within the monolayers in a function of time, depending on the experiment type (e) and on the cell type within the WT+RasV12 co-culture experiments (f), as shown in (a) and (b). Two-sample *t* -test: WT vs. RasV12, WT vs. Mix, and RasV12 vs. Mix, p *<* 0.001 (a); WT vs. RasV12, WT vs. Mix, and RasV12 vs. Mix, p *<* 0.001 (c); WT vs. RasV12, p *<* 0.001, WT vs. Mix, p *>* 0.1, RasV12 vs. Mix, p *<* 0.001 (e). Data is presented as the mean *±* standard deviation (s.d.).

In the analysis described above, the mechanical quantities were averaged over the whole monolayer without distinguishing between cell types. When we analyzed the data from the WT+RasV12 monolayer to differentiate between WT and RasV12 cells, both the traction and inter-cellular stress levels were closely aligned between the two cell types (green and magenta lines in Fig. 2b, d, f). These data clearly indicated that co-culturing WT and RasV12 cells induced a shift in traction and intercellular stress, leading them to converge to comparable levels.

### 2.2 Calcium sparks in WT cells did not alter traction and intercellular stress in neighboring cells

A decrease in isotropic intercellular stress in WT cells induces transient upsurges of intracellular calcium, called calcium sparks [33]. To address the potential changes in cell mechanics following calcium sparks, we tracked the traction and inter-cellular stress over time in regions proximal to the cells producing calcium sparks. Fig. 3 shows no significant trend in the evolution of any mechanical quantity in the area surrounding the cells undergoing a calcium spark event (ROI radius: d = 53 μm in Fig. 3, d = 29 μm and d = 102 μm in Fig. A2). These results exclude the possibility of a feedback mechanism from calcium sparks to the traction and intercellular stress.

**Fig. 3.**
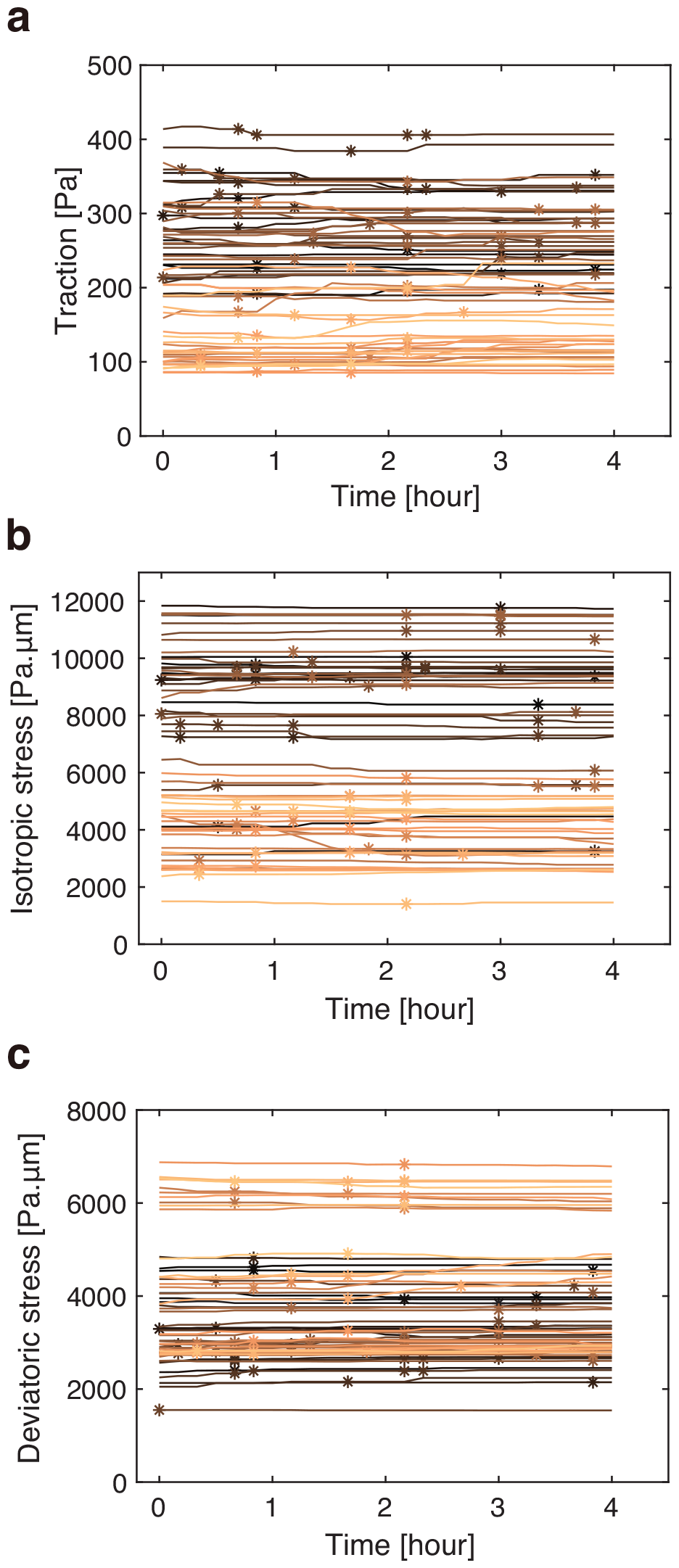
Traction and intercellular stress remain unchanged before and after calcium sparks in MDCK cells. (a–c) Quantification of traction (a), isotropic stress (b), and deviatoric stress (c) values near cells exhibiting calcium sparks in the WT+RasV12 monolayer comprising GCaMP WT MDCK cells and Myc-tagged RasV12-induced MDCK cells. The horizontal axis indicates time after the start of image acquisition, with the 0-hour mark corresponding to 6 h after induction of RasV12 expression. Measurements were made within a 53 μm radius surrounding cells exhibiting calcium sparks. Lines represent data for each ROI. Calcium spark occurrences are indicated by stars.

### 2.3 Convergence of traction and intercellular stress in co-culture of other cell types

Next, we investigated whether the convergence of traction and intercellular stress upon co-culturing different cell lines can be observed in other cell combinations. For this, we turned to MDCK-II cells, which are known to exhibit a more spread morphology and higher motility than MDCK cells [38]. Mechanical measurements were conducted as in MDCK cells, except that MDCK-II cells adhered to soft substrates through fibronectin and were confined in 1.8 mm diameter circles (Fig. 1e-h; Materials and Methods) [39]. To eliminate the potential effects of cell proliferation, such as density-dependent jamming [40], we treated the cells with a cell cycle inhibitor, Mitomycin C, before observation (Materials and Methods).

MDCK-II cells with different cell-cell adhesion strengths were examined in this study. Such cell-cell adhesion differences can lead to segregation into domains of distinct cell types [15, 16, 41]. Specifically, we used E-cadherin (E-cad) knockout (KO) cells [42]. The E-cad KO cells maintain a cohesive monolayer through Cadherin-6 [43].

TFM analysis confirmed that the traction exerted by the E-cad KO cells was higher than that exerted by the WT cells during the early phase (Fig. 4a), which is consistent with previous studies showing that E-cad KO results in the higher traction and an increase in the size of the focal adhesions [43]. As expected from the elevated traction, the E-cad KO monolayer displayed larger isotropic and deviatoric stresses than the WT monolayer (Fig. 4a, c, e; Video A2). When the WT and E-cad KO cell were co-cultured, the traction and intercellular stress dropped below those observed in the WT monolayer (Fig. 4a, c, e; Video A2). Within the mixed monolayer, both the WT and E-cad KO cells exhibited traction and inter-cellular stress of the same magnitude, respectively (Fig. 4b, d, f). Similar to the case of cell competition, the size of the ROI did not impact the evolution of mechanical quantities (Fig. A1b, d, f, h). Collectively, these findings suggest that the convergence of traction and intercellular stress in mixed monolayers is a general characteristic of epithelial cell populations and is not specific to the context of cell competition.

**Fig. 4.**
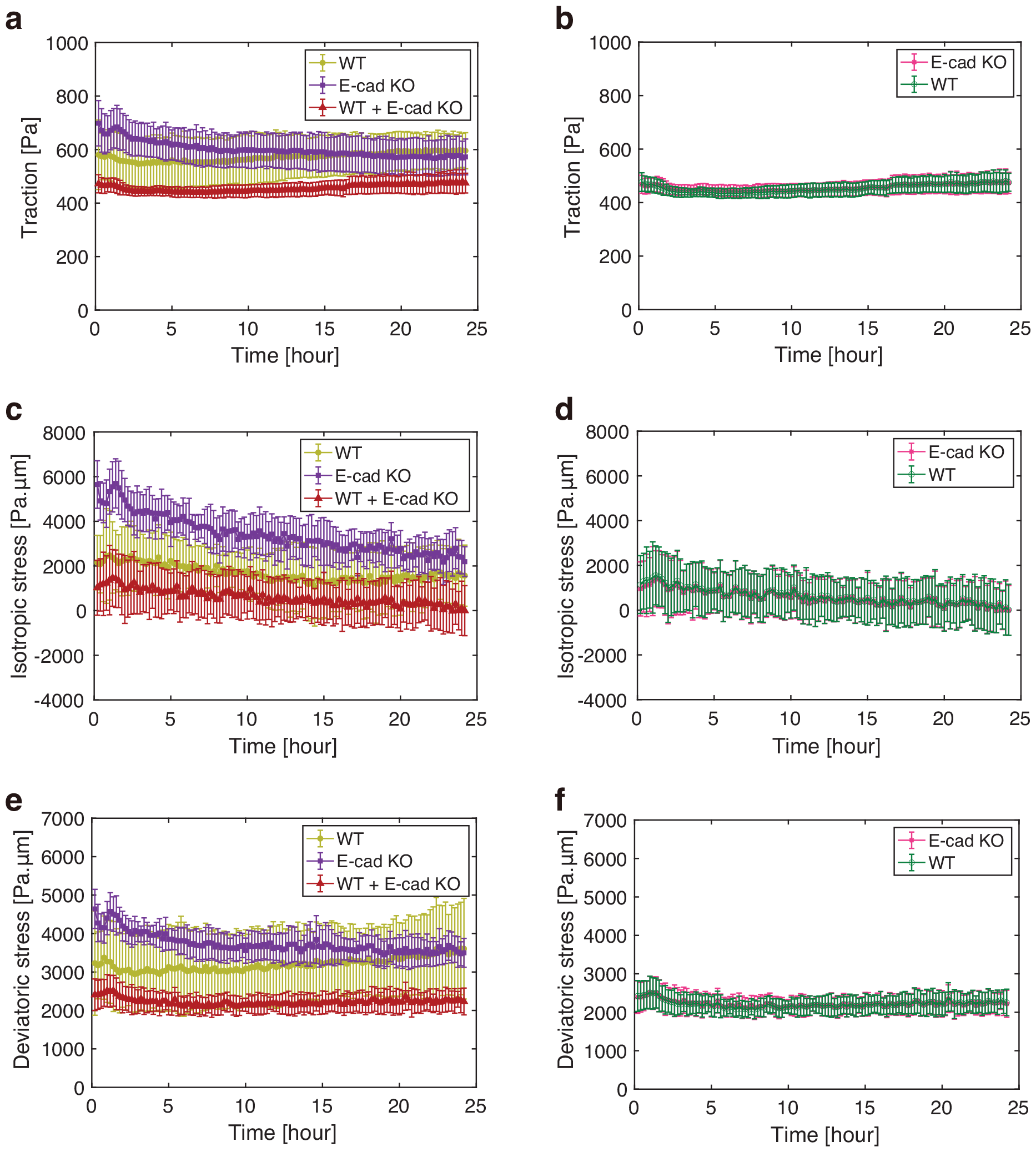
Quantification of traction, isotropic stress and deviatoric stress upon mixing WT and E-cad KO MDCK-II cells. (a, b) Magnitude of the traction exerted by the cells in a function of time, depending on the experiment types (a) and on the cell types within the WT+ E-cad KO co-culture experiments (b) (WT: n = 8, E-cad KO: n = 6, and WT+ E-cad KO: n = 8). 0 hour corresponds to 24 h after the cell seeding. In (a), WT represents the WT MDCK-II cells in single-cell colonies (light green), E-cad KO represents the E-cad KO MDCK-II cells in single-cell colonies (purple), and WT+E-cad KO represents the co-culture of WT and E-cad KO MDCK-II cells (red). In (b), the traction magnitude of each cell type (WT: green, E-cad KO: magenta) in the WT+E-cad KO co-culture experiments is plotted. (c, d) Isotropic stress within the monolayers in a function of time, depending on the experiment type (c) and on the cell type within the WT+E-cad KO co-culture experiments (d), as shown in (a) and (b). (e, f) Magnitude of deviatoric stress within the monolayers in a function of time, depending on the experiment type (e) and on the cell type within the WT+E-cad KO co-culture experiments (f), as shown in (a) and (b). Two-sample *t* -test: WT vs. E-cad KO, WT vs. Mix, and E-cad KO vs. Mix, p *<* 0.001 (a); WT vs. E-cad KO, WT vs. Mix, and E-cad KO vs. Mix, p *<* 0.001 (c); WT vs. E-cad KO, WT vs. Mix, and E-cad KO vs. Mix, p *<* 0.001 (e). Data is presented as the mean *±* s.d.

### 2.4 Intercellular stress exhibited no strong dependency on the distance to the cell type boundaries in co-culture experiments

To identify factors contributing to the convergence of intercellular stress upon mixing different cell types, we investigated whether the proximity of one cell type had any effect on the intercellular stress within the other cell types. To this end, we analyzed the correlation between intercellular stress and the shortest distance to cell type boundaries. We observed no clear dependence on the distance to the cell type boundaries for either isotropic stress or deviatoric stress magnitude at 8 h after the induction of RasV12 expression in the WT and RasV12 co-culture (Fig. 5a, c), and at 42 h after the cell seeding in the WT and Ecad KO co-culture, respectively (Fig. 5b, d). We also determined that the distance dependency was negligible at earlier time points. In the WT and RasV12 co-cultures, the absolute values of the correlation between the stress magnitude and the distance to the cell type boundaries (|*R*^2^|) were *<* 0.02 at 6 h after the induction of RasV12 expression. In the WT and Ecad KO co-culture, |*R*^2^| was *<* 0.03 at 24 h after the cell seeding. These results show that the change in intercellular stress is not restricted to the cell type boundaries, but instead suggests the possibility that it is collectively induced within the monolayer through a mechanism that can act over long distances.

**Fig. 5.**
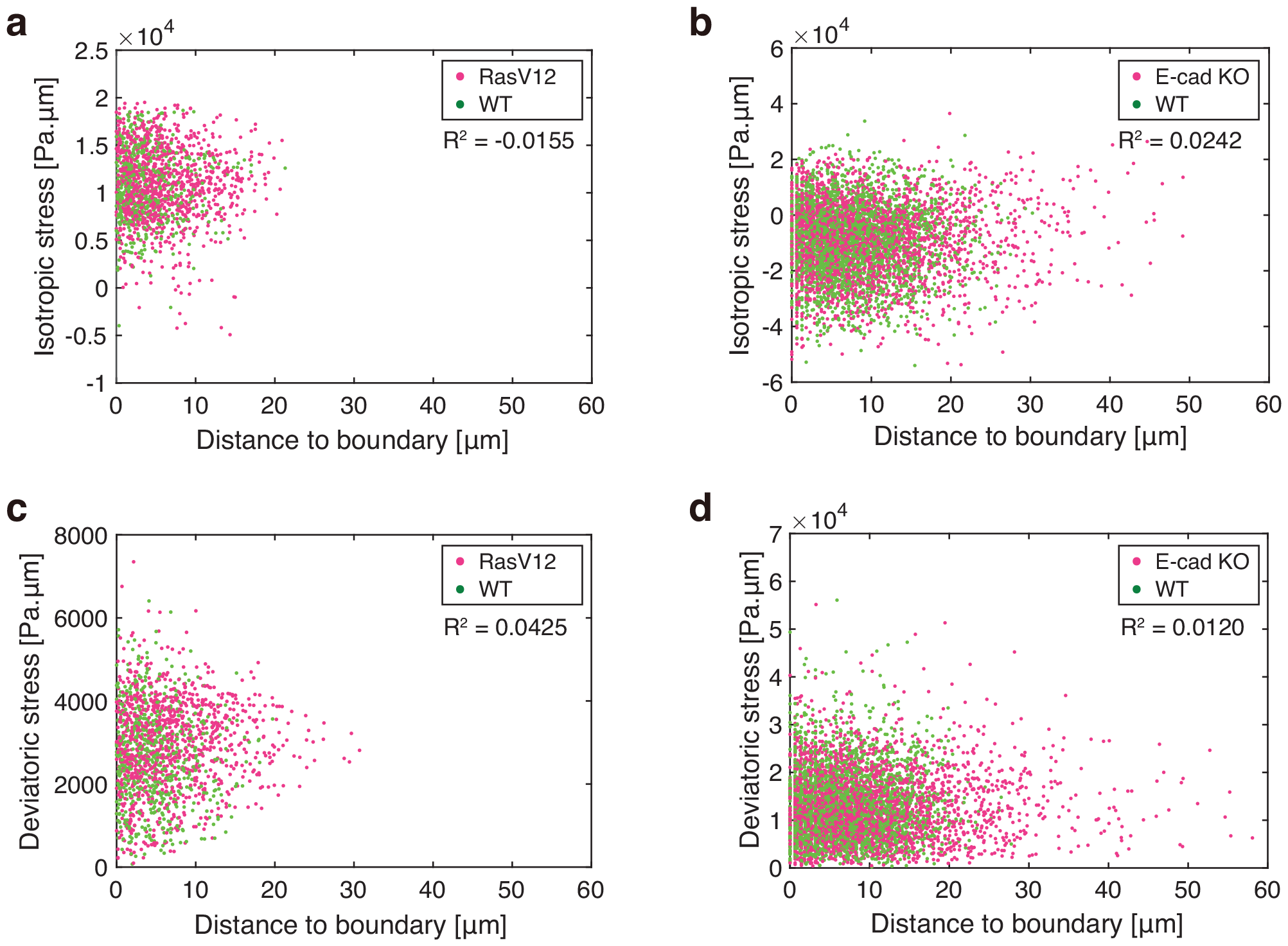
Dependency of intercellular stress magnitude on the distance from the cell type boundaries in co-culture experiments. (a–d) Scatter plots of isotropic stress (a, b) and deviatoric stress magnitude (c, d) in co-culture experiments in a function of the distance to the closest boundary between the two mixed cell types within the monolayer. The correlation coefficient is shown in the top-right corner. (a, c) Data obtained at 8 h after the induction of RasV12 expression are plotted. Green and magenta represent WT MDCK cells and RasV12 MDCK cells, respectively. (b, d) Data obtained at 42 h after the cell seeding (18 h after the start of image acquisition) are plotted. Green and magenta represent WT MDCK-II cells and E-cad KO MDCK-II cells, respectively.

### 2.5 Continuum model for the self-propelled binary cell mixture identifies physical conditions for mechanical convergence

To elucidate a long-range mechanism driving the mechanical convergence in mixed-cell populations, we introduced a two-dimensional hydrodynamic model for a mixture of two types of active cells [44–47]. In our model, different types of cells inherently have different viscosities and self-propelled forces. This formulation of self-propelled cell populations is motivated by experimental observations showing that different cell types in mixed-cultures moved at the same average speed and in the same orientation, despite exhibiting distinct velocities when cultured separately, a phenomenon particularly evident in MDCK-II cells, which exhibited higher motility than MDCK cells (Fig. 6). In addition, to reflect long-range rotational movement in the MDCK-II monolayer, we introduced a polar variable representing cellular polarity, along which cells move.

**Fig. 6.**
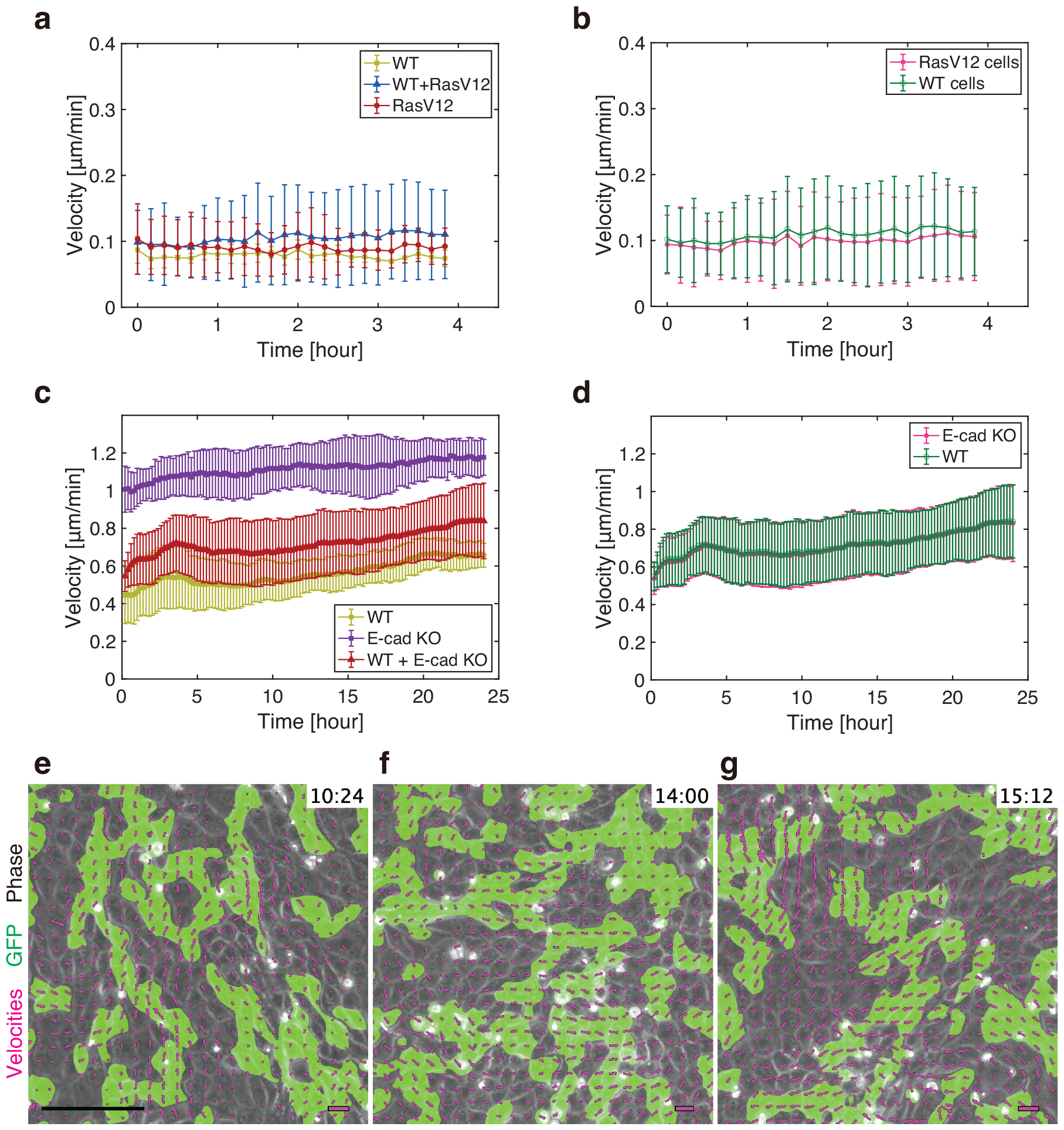
Quantification of velocity magnitudes and orientations in co-culture experiments. (a, b) Magnitude of velocity during the early phase of cell competition in within MDCK cell monolayers in a function of time, depending on the experiment types (a) and on the cell types within the WT+RasV12 co-culture experiments (b) (n = 3 for each plot). 0 hour corresponds to 6 h after the induction of RasV12 expression. In (a), WT represents the co-culture of GCaMP WT and non-tagged WT MDCK cells (light green), WT+RasV12 represents the co-culture of GCaMP WT and Myc-tagged RasV12-induced MDCK cells (blue), and RasV12 represents the co-culture of CMFDA-stained RasV12-induced and unstained RasV12 MDCK cells (red). In (b), the velocity magnitude of each cell type (WT: green, RasV12: magenta) in the WT+RasV12 co-culture experiments is plotted. (c, d) Magnitude of velocity upon mixing WT and E-cad KO MDCK-II cells in a function of time, depending on the experiment type (c) and on the cell type within the WT+E-cad KO co-culture experiments (d) (WT: n = 8, E-cad KO: n = 6, WT+ E-cad KO: n = 8). 0 hour corresponds to 24 h after the cell seeding. In (c), WT represents the WT MDCK-II cells in single-cell colonies (light green), E-cad KO represents the E-cad KO MDCK-II cells in single-cell colonies (purple), and WT+E-cad KO represents the co-culture of WT and E-cad KO MDCK-II cells (red). In (d), the velocity magnitude of each cell type (WT: green, E-cad KO: magenta) in the WT+E-cad KO co-culture experiments is plotted. (e–g) Phase contrast (gray), GFP mask (green) and velocity field (magenta arrows) at the indicated time in mixed-culture of GFP WT MDCK-II cells and non-tagged E-cad KO MDCK-II cells. To allow for better readability, velocity arrows are scaled to match the grid size with the maximum velocity value at each frame being equal to one unit of grid size. Data in (a–d) is presented as the mean *±* s.d. Scale bars: 150 μm (e, lower left), 2 μm/min (e-g, lower right)

We extended the active polar model into a binary mixture, employing an approach similar to that used for the active nematic model in [45, 47]. The model is described by a phase-field variable *ϕ*(***r***, *t*), cellular polarity ***p***(***r***, *t*), and velocity ***v***(***r***, *t*), where ***r*** and *t* represent position and time, respectively. The phase field *ϕ* is introduced to represent the binary cell mixture, where *ϕ* = 1 indicates the area occupied by cell type A, and *ϕ* = −1 denotes the area occupied by cell type B. Free energy associated with these fields is introduced in the generic form as

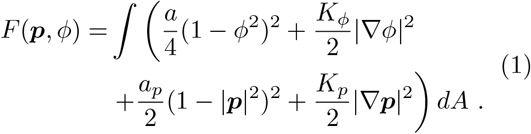

The time evolution of these fields is given by

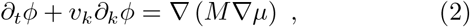

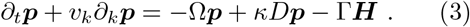

Here, *μ* = *δF/δϕ* and ***H*** = *δF/δ****p*** represent the the chemical potential for *ϕ* and molecular field for ***p***, respectively (Materials and Methods). *D* ≡ (**∇***v* + **∇***v*^*T*^)*/*2 and Ω ≡ (**∇***v* − **∇***v*^*T*^)*/*2 are the symmetric and anti-symmetric parts of the velocity gradient tensor. *κ* is a parameter for controlling the alignment of the cell polarity under a simple shear flow; we put *κ >* 0, indicating ‘rod-like’ in terms of liquid crystal theory, where the cell polarity is enhanced in the direction of shear extension. Parameters *M* and Γ are parameters to determine the time scale of cell mobility and the cell polarity dynamics.

The cell populations are modeled as viscous Newton fluids, and the velocity field ***v***(*t*, ***r***) is determined by the force balance (Stokes) equation and the incompressible condition. We assumed incompressibility due to the absence of cell division.

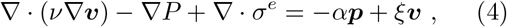

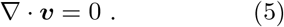

Here, *P* represents pressure, and elastic stress tensor *σ*^*e*^ is derived from the variational of the Free energy Eq. 1 (Materials and Methods). Cells interact with their substrate via a self-propelled active force *α****p*** and friction −*ξ****v***. In our system, the self-propelled force is the only active force driving the dynamics. As mentioned above, different types of cell populations exhibit distinct velocities, the coefficient of the self-propelled force is set as *α* = *α*_*A*_ for A cells and *α* = *α*_*B*_ for B cells, and thus the intrinsic cell migration speed is given as *α*_*A*_*ξ*^−1^ and *α*_*B*_*ξ*^−1^, respectively. The viscosities of the cells are also different, with the viscous coefficient *ν* = *ν*_*A*_ for A cells and *ν* = *ν*_*B*_ for B cells. Thus we took

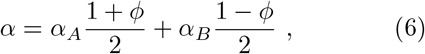

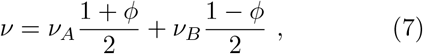

where *α*_*A*_ and *α*_*B*_ indicates self-propelled force coefficients of cell type A and B, respectively, etc. We set *α*_*A*_ = 2*α*_*B*_ and *ν*_*A*_ *< ν*_*B*_ assuming that A cells are faster and less viscous than B cells.

We numerically solved Eqs. 2–5 on a 50 *×* 50 square region. In the simulation, we set *α*_*A*_ = 1.0, *α*_*B*_ = 0.5, *ν*_*A*_ = 5.0, *ν*_*B*_ = 10.0, and *ξ* = 1.0. Then the intrinsic cell migration speed of cell A is two times faster than that of cell B, reflecting the experimental observation in MDCK-II cells (Fig. 6) (see Materials and Methods for the rationale of the parameter choice). The following parameters were also fixed: *a* = 0.1, *K*_*ϕ*_ = 0.05, Γ = 5.0, *M* = 10.0, *a*_*p*_ = 0.1, and *κ* = 0.7. To see the role of cell polarity alignment, we changed *K*_*p*_.

Fig. 7a, along with Video A3, illustrates an example from our numerical simulation that began with a random initial condition. In the mixture of the two phases, which inherently have a two-fold difference in intrinsic velocity, the mean velocities converged to comparable levels (solid lines in Fig. 7d). Similarly, the stress magnitudes became indistinguishable among the two phases (solid lines in Fig. 7e). In a different set of simulations, reducing the coupling of cell polarity led to a more disordered pattern in the velocity field, yet the significant shift in the mean velocity and the isotropic stress was still apparent upon mixing (Fig. 7b and Video A4; dashed lines in Fig. 7d, e). These results demonstrate that the global shift and convergence in the stress and velocity fields can be collectively induced within the mixed-cell population.

**Fig. 7.**
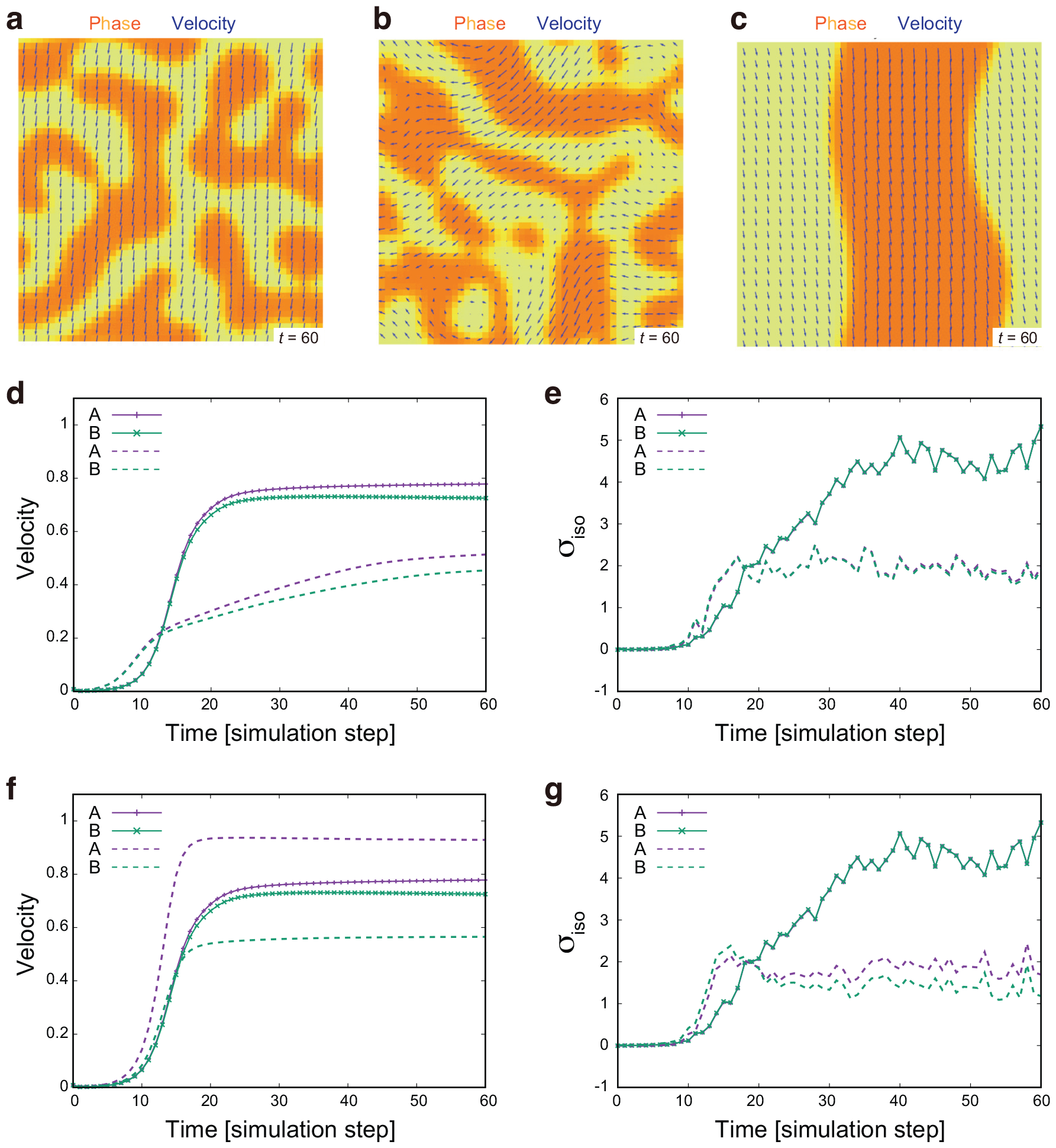
Numerical simulation of binary cell mixture and resulting velocity and stress profiles. (a–c) Visualization of the phase field (color scale) and velocity field (blue arrows) at the indicated simulation time. Orange and yellow indicate regions occupied by cells A (*ϕ* = 1) and cells B (*ϕ* = −1), respectively. Parameters for the self-propelled force and viscosity are set as *α*_*A*_ = 1.0, *α*_*B*_ = 0.5, *ν*_*A*_ = 5.0, and *ν*_*B*_ = 10.0 (see the main text for the definition of parameters). (a) Simulation started from a random initial condition with the polarity alignment parameter set to *K*_*p*_ = 1.0. See Video A3 for the full sequence. (b) Simulation started from the random initial condition with *K*_*p*_ = 2.0 *×* 10^−2^. See Video A4 for the full sequence. (c) Simulation conducted with the same parameter set as in (a), but with a different initial condition that completely segregates the two phases. See Video A5 for the full sequence. (d, e) Temporal evolution of the mean velocity (d) and 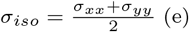 for individual cell types (cells A: purple, cells B: green), with solid lines in corresponding to the data from (a) and dashed lines to (b). Note that purple and green solid lines overlapped in (e). (f, g) Temporal evolution of the mean velocity (f) and 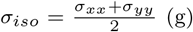 for individual cell types (cells A: purple, cells B: green), with solid lines in corresponding to the data from (a) and dashed lines to (c). In these simulations, the parameters are set as *η* = 1.0, *a* = 0.1, *K*_*ϕ*_ = 0.05, Γ = 5.0, *M* = 10.0, *a*_*p*_ = 0.1, *ξ* = 0.7.

Moreover, our numerical simulations revealed that the extent of mechanical convergence was influenced by the degree of phase separation. Using the same parameters as in Fig. 7a but starting from an initial condition of completely segregated phases, each phase retained its distinct velocities and stresses (Fig. 7c and Video A5; dashed lines in Fig. 7f, g). This data suggests that the mechanical behavior of cells is significantly affected by the spatial arrangement of the different cell types in mixed-cell populations.

## 3 Discussion

In this study, we determined that co-culture with different cell types can significantly alter the traction, intercellular stress, and velocity within a monolayer of mammalian epithelial cells. Our findings suggest that interactions with neighboring cells are pivotal in determining mechanical behaviors of cells. This challenges the common assumption that cell mechanics, being largely genetically determined, remain consistent irrespective of interactions with other cell types. Comparable shifts in cell mechanics have been observed at the interface between distinct cell types [48–50]; for example, junction tension increases at compartment boundaries in embryos and at clonal boundaries separating normal and oncogenic cells. Our *in situ* mechanical measurements revealed that the mechanical changes were not localized, but spread throughout the entire monolayer without dependence on the distance to the cell type boundaries. These observations highlight a collective mechanism that acts over long distances for triggering the mechanical convergence. Investigating the distribution and activity of molecules that control cell mechanics is crucial to further support our findings and shed light on the mechanisms by which interactions between diverse cell types are integrated with genetical regulations to control the mechanics of mixed-cell populations.

The mechanisms underlying the differences between localized changes at the interfaces and more global changes within the entire monolayer remain elusive. One critical factor could be the cell mixing ratio, which was set to 1:1 in this study. Determining the specific cell mixing ratio at which a mixed monolayer begins to behave as a cohesive entity is of interest. In addition, the interface geometry should be considered because it can influence both the local cell mixing ratio and the magnitude of intercellular signaling [51, 52] (see also discussion about the modeling results). Biological tissues may utilize these geometrical parameters to dynamically adjust cell mechanics during tissue development and repair.

Differential cell adhesion has been known to induce cell sorting [15, 16, 41]. However, in our experiments with WT and E-cad KO cells in MDCK-II monolayers, no significant segregation into large domains were observed. To eliminate the potential effects of cell proliferation and jamming, we incubated the cells with a cell cycle inhibitor for 1.5 h. This inhibition of cell proliferation prevented clonal expansion, which may have contributed to domain expansion among cells of the same type. Without cell proliferation, additional mechanisms, such as ephrin-Eph-based repulsion [53], may be required for the efficient sorting of cells in MDCK-II monolayers.

We introduced a simple hydrodynamic model for a binary cell mixture and uncovered that two phases, each with intrinsically distinct self-propelled forces, can behave like a single entity when mixed. We speculate that the force transmission within the cell population aligns the movement of the two phases, leading to a global alteration in the stress field. The well-segregation of phases however impedes this coherent movement, thus preventing mechanical convergence. As long as the cellular domains remain small, a moderate level of disorder in the polarity and velocity fields has a minor impact on the mechanical convergence in mixed-cell populations. The insights gleaned from this model provide clues to a cell-cell communication mechanism that drives the observed global changes in velocity and stress fields in co-culture experiments.

Although our mixed-phase model suggests a physical mechanism driving the convergence of traction, intercellular stress, and velocity, it does not account for the direction of the shifts in these mechanical quantities. In the WT+RasV12 monolayer, we observed that the values of traction and intercellular stress were intermediate to those in the WT and RasV12 monolayers, while the velocity exceeded that seen in single-cell experiments. Conversely, in the WT+E-cad KO monolayer, the velocity fell between those in the WT and E-cad KO monolayers, yet the traction and intercellular stress were lower than those in single-cell experiments. These variations underscore the complex interactions between different cell types and highlight the need for further investigation to fully understand the underlying dynamics governing these mechanical shifts.

In conclusion, this study highlights the importance of *in situ* mechanical measurements in mixed-cell populations for understanding the mechanics of multicellular systems. As the available mechanical tools continue to expand [25, 26], we anticipate that various mechanical quantities will be directly quantified in mixed-cell populations. This will enable future studies to explore the intricate interplay between the genetics and mechanics in heterotypic tissue environments.

## 4 Materials and Methods

### 4.1 Cell lines

MDCK-II and MDCK cells were used in this study. WT MDCK-II cells and E-cad KO MDCK-II cells were a gift from Tetsuhisa Otani. WT-GFP MDCK-II cells were generated by transfecting MDCK II cells with pCANw-EGFP (CAG-GFP-IRES-neo) [54] using Lipofectamine LTX followed by selection with 400 μg/ml G418. The E-cad KO MDCK-II cell line was established in [42]. The MDCK cell lines used are described in [33].

### 4.2 Cell culture

MDCK-II cells were cultured in Dulbecco’s modified Eagle’s medium (DMEM) (Gibco) supplemented with 10% fetal bovine serum (FBS) (Biowest) and 1% penicillin/streptomycin (Gibco) at 37°C and 5% CO_2_. WT-GFP MDCK-II cells were maintained with 500 μg/ml of G418 (Geneticin) (Gibco). MDCK cells were cultured in DMEM (Wako) supplemented with 10% FBS (Sigma-Aldrich), 1% penicillin/streptomycin (Life Technologies), and 1% GlutaMax (Life Technologies) at 37°C and 5% CO_2_. MDCK-GCaMP6s cells and MDCK-pTRE3G Myc-RasV12 cells were maintained with 80 μg/ml of G418 (Geneticin) (Gibco) and 0.5 μg/ml blasticidin (Invitrogen), respectively [55, 56]. Mycoplasma contamination was regularly tested in all cell lines using a MycoAlert Mycoplasma Detection Kit (Lonza).

### 4.3 Preparation of soft substrate

Experiments were performed in a 35-mm glass-bottom dish (Iwaki). The glass bottom of each dish was covered with a soft gel created by mixing parts A and B of the silicone elastomer DOWSIL CY 52-276 (Dow Corning). For experiments with MDCK-II cells, a soft substrate with a Young’s modulus of 15 kPa was obtained by mixing parts A and B in a 1:1 ratio, whereas a soft substrate with a Young’s modulus of 3 kPa (A:B ratio of 6:5) was used for experiments with MDCK cells [57]. Soft substrate dishes were then covered with 200-nm diameter, fluorescent beads (red for MDCK-II experiments and dark red for MDCK experiments) (Invitrogen) to track the traction exerted by the cells over time.

### 4.4 Live cell imaging

In the experiments with MDCK-II cells, the cells were treated with Mitomycin C (Sigma-Aldrich), a cell cycle inhibitor, at a concentration of 10 μg/ml for 1.5 h, 24 h before cell seeding [58]. On the day of cell seeding, a microprinting technique was performed to stamp fibronectin (Wako) onto a soft substrate covered with beads [39]. The fibronectin was used at a concentration of 50 μg/ml and was stamped in the shape of 1.8 mm diameter circles. Pluronics F-127 (Sigma-Aldrich) at a concentration of 2% was used to prevent the cells from adhering outside of the circles. After 1 h of incubation, Pluronic F-127 was rinsed with PBS (Gibco) and MDCK-II culture medium. In experiments involving single-cell type cultures, cells were directly plated on a stamped soft substrate-covered dish, whereas in the case of mixed-cell type experiments, cells from each cell line were first transferred to a vial in the desired proportion and then seeded on the dish. After incubating the cells at 37°C with 5% CO_2_ for 1 h, the cells were rinsed with PBS and culture media until no floating cells remained. Image acquisition began 24 h later, following a change of the culture media. Images were captured using an inverted confocal spinning disk microscope (Olympus IX83 combined with Yokogawa CSU-W1) equipped with an iXon3 888 EMCCD camera (Andor), an Olympus 20x/NA0.45 LUCPLFLN PH dry objective, and a temperature control chamber (TOKAI HIT), using IQ 2.9.1 software (Andor). A number of 3x3 fields of view are required to capture a monolayer of cells on a 1.8 mm diameter circle. Images were captured at 37°C and 5% CO_2_ for 24 h at 12 min intervals.

Soft substrate dishes used for experiments with MDCK cells were coated with 0.3 mg/ml collagen at 4°C for 16 h and subsequently washed with PBS [33]. MDCK cells were incubated in Leibovitz’s medium (L-15) (Gibco) supplemented with 10% FBS. MDCK-GCaMP6s cells were mix-cultured with MDCK or MDCK-pTRE3G Myc-RasV12 cells at a ratio of 1:1 and plated on a soft substrate dish. To prepare a sample of RasV12 alone, MDCK-pTRE3G Myc-RasV12 cells were pre-stained with CellTracker Dye CMFDA (green) (Life Technology) before mixing with unstained MDCK-pTRE3G Myc-RasV12 cells. The mixture of cells was incubated for 12–16 h until a monolayer was formed, followed by the treatment of 1 μg/ml doxycycline (Sigma-Aldrich) for 6 h to induce the expression of RasV12. Images were captured using an inverted confocal microscope (Nikon A1 HD25) equipped with a Nikon 25x/NA1.05 PLAN APO silicon oil-immersion objective and a temperature control chamber (TOKAI HIT). Images were captured at 37°C for 4 h at 10 min intervals, starting at 6 h after the induction of RasV12.

### 4.5 Image analysis

#### Movie creation

For experiments with MDCK-II cells, images were re-ordered using ImageJ/Fiji to construct individual movies of phase contrast, GFP, and red fluorescent beads. For phase contrast and bead movies, the focused slice at each time frame was selected, whereas the z-slices of the GFP movies were projected to increase the signal-to-noise ratio. Images at different x- and y-positions were stitched together using the Fiji Grid/Collection stitching plugin [59] to cover 1.8-mm diameter circles. Finally, for both experiments with MDCK and MDCK-II cells, the bead movies were stabilized to remove any shifts in the x- and y-directions.

#### Mask creation

A custom-made ImageJ/Fiji macro was used to create black and white masks to isolate the monolayer of cells from the outside of the monolayer (made from the phase contrast movies and called cell masks), and to separate the two populations of cells within a mixed-culture experiment using the GFP signal of one of the cell lines (made from the GFP fluorescence movies and called GFP masks) (Fig. 1). We also employed black and white masks to calculate mechanical values within smaller ROIs, thereby excluding any values outside the chosen area from the calculation.

#### Velocity fields

Velocity fields were calculated using the MatPIV toolbox for MATLAB (Mathworks). This allowed us to perform Particle Image Velocimetry (PIV) on the phase-contrast movies between consecutive frames using interrogation window sizes of approximately 20 μm for MDCK cells experiments, and 40 μm for MDCK-II cells experiments, with 50% overlap in both cases.

#### Quantification of calcium sparks

MorpholibJ, a Fiji plugin, was used to segment the inverted bright field images and label the outer contours of individual MDCK cells. If necessary, the segmented images were corrected manually. In addition, a GFP mask was generated to identify cells expressing GCaMP6s. Each GFP-positive cell was tracked over time using the Trackmate plugin in Fiji. For the analysis, we excluded cells located at the edge of the field of view and those that could not be tracked over the entire duration of the experiment. Calcium sparks were defined as events in which the intensity of GCaMP in a cell exceeded the mean GFP intensity by more than five standard deviations [33]. Physical quantities (traction and isotropic and deviatoric stress values) were then averaged at each time frame in a circle of a chosen radius around the centroid of the selected cells and plotted over time for each cell (Fig. 3 and Fig. A2).

### 4.6 Traction force microscopy (TFM) and intercellular stress calculation

#### Traction force microscopy

PIV was used to track the bead displacements with interrogation windows of 16 and 10 μm used for the experiments performed respectively with MDCK and MDCK-II cells, and an overlap of 75%. Subsequently, the ImageJ/Fiji FTTC plugin [60] with a regularization factor of 8e-11 was used to obtain the tractions 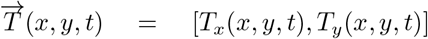 from the displacement fields.

#### Stress tensor calculation

Intercellular stress *σ* was obtained from the traction 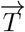 through the relationship div 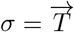 using Bayesian inversion stress microscopy (BISM) [27]. A two-dimensional symmetrical stress tensor 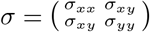 was calculated in each interrogation window of the PIV grid used for TFM. From each stress tensor *σ*, we calculated the isotropic stress *σ*_*iso*_ which is the trace of the tensor as 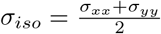. The value *σ*_*iso*_ is a scalar and the opposite of the pressure inside the tissue, so that *σ*_*iso*_ *<* 0 means compression in the tissue, and *σ*_*iso*_ *>* 0 means that the tissue is under tension. The deviatoric stress *σ*_*dev*_ is a symmetrical and traceless tensor defined as *σ*_*dev*_ = *σ*−*σ*_*iso*_*I* where *I* is the identity matrix. By applying a linear transformation to the deviatoric stress tensor *σ*_*dev*_, we determine its eigenvalues, *λ*_*dev*_ and −*λ*_*dev*_. Since *σ*_*dev*_ is a symmetrical and traceless tensor, these eigenvalues are equal in magnitude but opposite in sign. We use the absolute value |*λ*_*dev*_| to represent the magnitude of the deviatoric stress.

### 4.7 Statistics

P-values were calculated in MATLAB using *ttest2* (two-sample t-test) and *ranksum* (two-sided Wilcoxon rank-sum test).

### 4.8 Model

#### Detailed expression of elastic stress, chemical potential, and molecular field

Elastic stress tensor *σ*^*e*^ = *σ*^*p*^ + *σ*^*ϕ*^ is derived from the free energy as

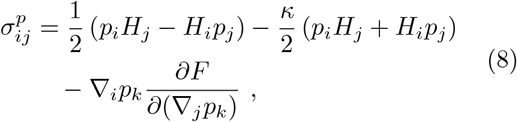

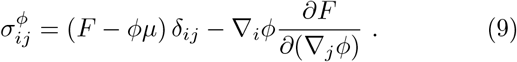

Detailed expressions for chemical potential *μ*, molecular field ***H***, and elastic stresses *σ*^*p*^ and *σ*^*ϕ*^ are as follows.

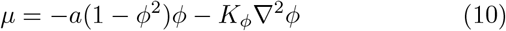

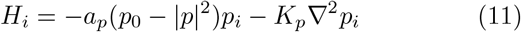

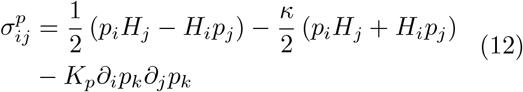

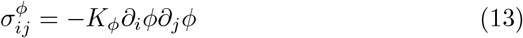

In general, the coefficients *ν, α, ξ*, and *K*_*p*_ can be functions of *ϕ*. In this study, we assume differences in self-propelled forces and viscosity between two types of cells (Eqs. 6 and 7), while the other coefficients are constant. For calculating *μ*, we neglected the contribution from the derivative of the coefficients.

#### Parameters

In our simulation, time and length units were chosen as 10 min and 10 *μ*m, respectively. The system size, a 50 *×* 50 square region in the simulation unit, is equivalent to 100 *×* 100 *μ*m^2^. The simulation duration in Fig. 7d-g corresponds to 600 min. Characteristic quantities were determined as follows. Intrinsic cell migrating velocities are *α*_*A*_*ξ*^−1^ = 1.0 *μ*m*/*min and *α*_*B*_*ξ*^−1^ = 0.5 *μ*m*/*min, reflecting the experimental observation in E-cad KO and WT MDCK-II cells (Fig. 6). *M*^−1^ = 1 min was set so that *ϕ* maintains the sharp interface during simulations. Γ^−1^ = 2 min indicates the characteristic time scale for cell polarity formation, while *ξ*^−1^ = 10 min does the characteristic time scale for velocity correlation. The interface width of the two-cell population is given as 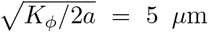. The correlation length for the polar field estimated by 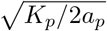 is ∼ 20 *μ*m at *K*_*p*_ = 1.0 (Fig. 7a, c) and ∼ 3 *μ*m at *K*_*p*_ = 0.02 (Fig. 7a, c). The correlation length of the velocity field is estimated by 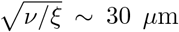. The temporal and spatial correlations of cell velocity are comparable with our experimental observations in MDCK-II cells. *κ* is a non-dimensional parameter and was set to *κ* = 0.7 following [45]. Collectively, these parameter values are within reasonable ranges.

#### Numerical simulations

We solved Eqs. 1-4 on a *L × L* = 50 *×* 50 square region with periodic boundary conditions. The square region is discretized by triangular mesh, whose size is at most 1.0. The simulation was carried out by a finite element method (FEM) using a FEM solver, FreeFEM++ [61]. Pressure is approximated using the P1 finite element, while other variables are by P2. For solving Eqs. 2 and 3, the Crank-Nicolson scheme was used, while the FreeFEM++ operator **convect()** was applied for calculating the convection term [61]. Time discretization was Δ*t* = 1.0 *×* 10^−2^.

## Acknowledgments

The authors would like to thank Tetsuhisa Otani for reagents, Yusuke Kishi, Kei Kozawa, and Benoit Ladoux for technical advice and support. This study was financially supported by JSPS KAKENHI Grant (17H05617) and the AMED PRIME program (20gm5810025h9904) to K.S., and JST CREST (JPMJCR1923) and JPJSBP (JPJSJPR 201915010) to S.I.

## Author contributions

E.G. and K.S. designed the research. E.G. and K.K. performed the experiments. K.I. assisted with the experiments. E.G. and T.N. analyzed the data. K.K. assisted with the data analysis. S.I. conducted numerical simulations. Y.F. supervised the project. E.G., K.S., and S.I. drafted the manuscript. All authors approved the final version of the manuscript.

## Data availability

The authors declare that the data supporting the findings of this study are available within the paper and its Supplementary files. The data are available from the lead contact upon reasonable request. The code used for numerical simulations can be downloaded from https://github.com/IshiharaLab/BinaryActivePolarModel.

## Declaration of Interests

The authors declare no competing interests.

## Appendix A Supplementary Figures

**Figure A1** We varied the size of the ROIs to assess potential size and positional dependencies in the measured mechanical quantities. Given that the patterns employed have a typical length *R* (either the radius of the circles or the side length of the square fields of view), we performed the quantification of traction, isotropic stress, and deviatoric stress in areas that have typical lengths of 0.95*R* and 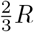. In both cases, the results closely resemble those obtained in the whole domains (compare Fig. A1 with Fig. 2 and Fig. 4).

**Figure A2** We tracked the traction and inter-cellular stress over time in regions proximal to the cells producing calcium sparks in Fig. 3. Here, we verified that the size of the area of calculation had no significant effect on the evolution of the mechanical quantities surrounding calcium spark events.

## Appendix B

### Video A1 Intercellular stress in monolayers of WT MDCK cells, RasV12-induced MDCK cells, and mixed-cultures of WT+RasV12 cells

Overlay of isotropic stress as colormaps with principal stress tensors as elliptical principal axes for WT MDCK cell monolayers (a), RasV12-induced MDCK cell monolayers (b), and mixed-cultures of WT and RasV12-induced cells (c). The stress tensor axes are presented in red when the stress value is positive, and in blue when it is negative. Scale bar: 100 μm.

### Video A2 Intercellular stress in monolayers of WT MDCK-II cells, E-cad KO MDCK-II cells, and mixed-cultures of WT+E-cad KO cells

Overlay of isotropic stress as colormaps with principal stress tensors as elliptical principal axes for WT MDCK-II cell monolayers (a), E-cad KO MDCK-II cell monolayers (b), and mixed-cultures of WT and E-cad KO cells (c). The stress tensor axes are presented in red when the stress value is positive, and in blue when it is negative. Scale bar: 500 μm.

### Video A3 Numerical simulation of binary cell mixture shown in Fig. 7a

The phase field (color scale) and velocity field (blue arrows) at the indicated simulation time. Orange and yellow indicate regions occupied by cells A (*ϕ* = 1) and cells B (*ϕ* = −1), respectively. Parameter values are described in the legend of Fig. 7.

### Video A4 Numerical simulation of binary cell mixture shown in Fig. 7b

The phase field (color scale) and velocity field (blue arrows) at the indicated simulation time. Orange and yellow indicate regions occupied by cells A (*ϕ* = 1) and cells B (*ϕ* = −1), respectively. Parameter values are described in the legend of Fig. 7.

### Video A5 Numerical simulation of binary cell mixture shown in Fig. 7c

The phase field (color scale) and velocity field (blue arrows) at the indicated simulation time. Orange and yellow indicate regions occupied by cells A (*ϕ* = 1) and cells B (*ϕ* = −1), respectively. Parameter values are described in the legend of Fig. 7.

**Fig. A1.**
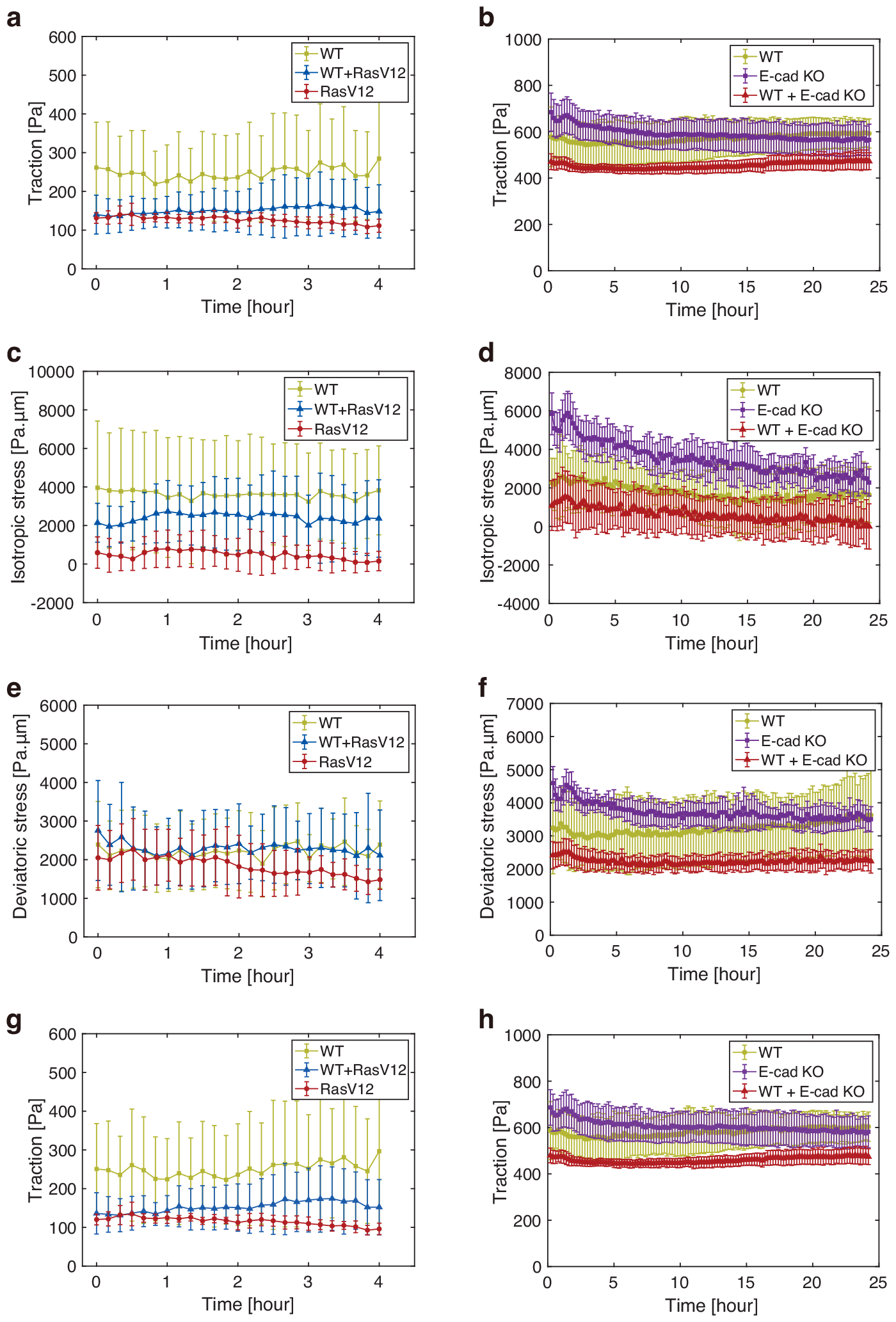
Quantification, in areas of typical length 0.95*R* and 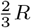, of traction, isotropic stress, and deviatoric stress. (a, b) Magnitude of the tractions, in areas of typical length 0.95*R*, exerted by the cells in a function of time depending on the experiment types in MDCK cells (n = 3 for each plot) (a) and in MDCK-II cells (WT: n = 8, E-cad KO: n = 6, and WT+ E-cad KO: n = 8) (b). In (a), 0 hour corresponds to 6 h after the induction of RasV12 expression. WT represents the co-culture of GCaMP WT and non-tagged WT MDCK cells (light green), WT+RasV12 represents the co-culture of GCaMP WT and Myc-tagged RasV12-induced MDCK cells (blue), and RasV12 represents the co-culture of CMFDA-stained RasV12-induced and unstained RasV12 MDCK cells (red). In (b), 0 hour corresponds to 24 h after the cell seeding. WT represents the WT MDCK-II cells in single-cell colonies (light green), E-cad KO represents the E-cad KO MDCK-II cells in single-cell colonies (purple), and WT+E-cad KO represents the co-culture of WT and E-cad KO MDCK-II cells (red). (c, d) Isotropic stress, in areas of typical length 0.95*R*, in a function of time depending on the experiment type in MDCK cells (c) or MDCK-II cells (d), as shown respectively in (a) and (b). (e, f) Magnitude of deviatoric stress, in areas of typical length 0.95*R*, in a function of time depending on the experiment type in MDCK cells (e) or MDCK-II cells (f), as shown respectively in (a) and (b). (g, h) Magnitude of the tractions, in areas of typical length 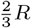, exerted by the cells in a function of time depending on the experiment types in MDCK cells (f) or MDCK-II cells (g), as shown respectively in (a) and (b). Data is presented as the mean *±* s.d.

**Fig. A2.**
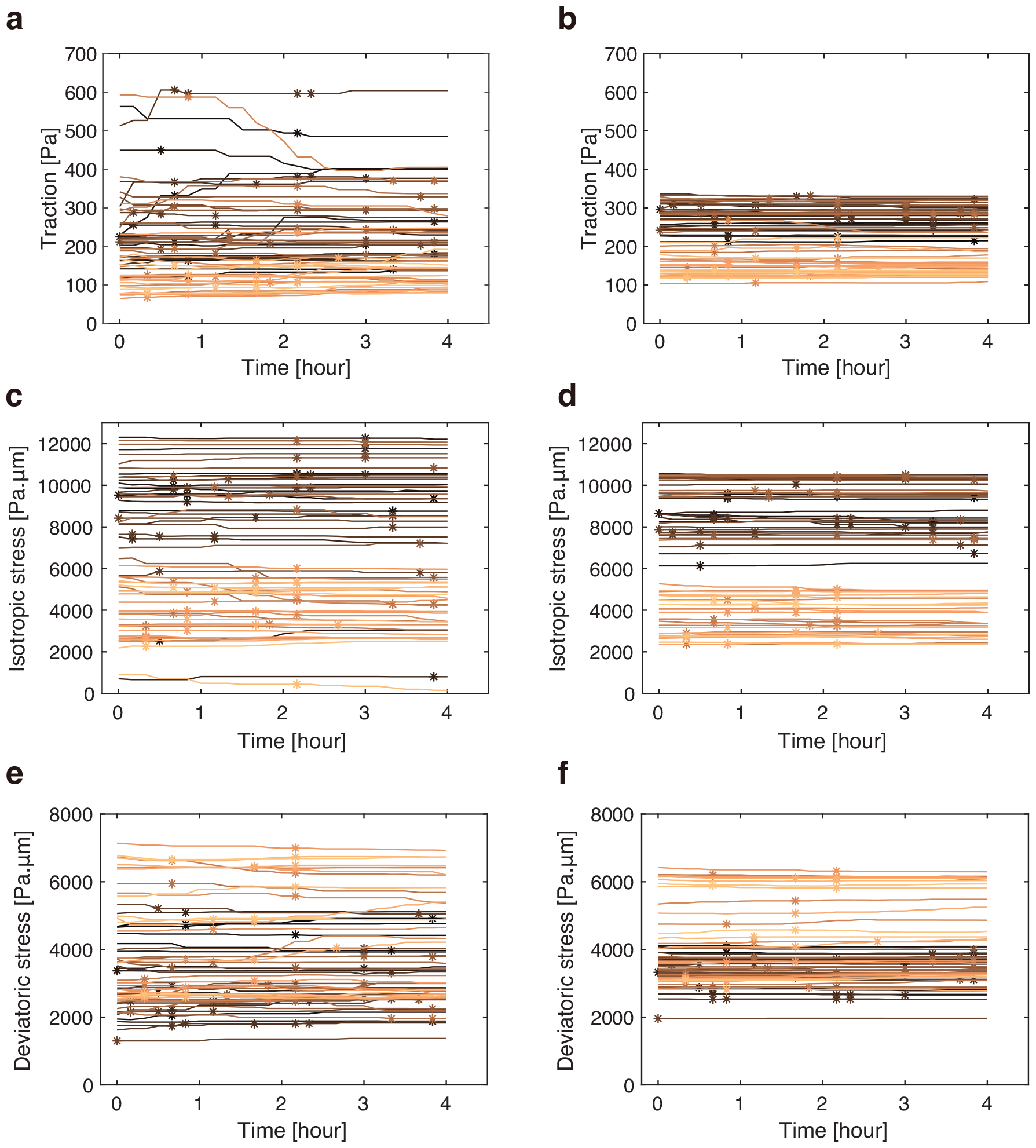
Evolution of mechanical quantities calculated in areas of different sizes surrounding calcium spark events. Quantification of traction (a, b), isotropic stress (c, d), and deviatoric stress (e, f) values in an area around cells producing a spark over the time of the experiment. In (a), (c), and (e), the radius of the area around the sparking cells used to calculate the averaged quantities is 29 μm; and in (b), (d), and (f), the radius is 102 μm. The horizontal axis indicates time after the start of image acquisition, with the 0-hour mark corresponding to 6 h after induction of RasV12 expression. Lines represent data for each ROI, and calcium spark occurrences are indicated by stars.

## Notes

### Competing Interest Statement

The authors have declared no competing interest.

## References

[1] J. B. Gurdon, E. Tiller, J. Roberts, and K. Kato. A community effect in muscle development. Current Biology, 3(1):1–11, 1993.

[2] Chii J. Chan, Carl-Philipp Heisenberg, and Takashi Hiiragi. Coordination of Morphogenesis and Cell-Fate Specification in Development. Current Biology, 27(18):R1024–R1035, 2017.

[3] Pulin Li and Michael B. Elowitz. Communication codes in developmental signaling pathways. Development, 146(12):dev170977, 2019.

[4] Yoshiki Sasai. Cytosystems dynamics in self-organization of tissue architecture. Nature, 493(7432):318–326, 2013.

[5] J. Gray Camp, Keisuke Sekine, Tobias Gerber, Henry Loeffler-Wirth, Hans Binder, Malgorzata Gac, Sabina Kanton, Jorge Kageyama, Georg Damm, Daniel Seehofer, Lenka Belicova, Marc Bickle, Rico Barsacchi, Ryo Okuda, Emi Yoshizawa, Masaki Kimura, Hiroaki Ayabe, Hideki Taniguchi, Takanori Takebe, and Barbara Treutlein. Multilineage communication regulates human liver bud development from pluripotency. Nature, 546(7659):533–538, 2017.

[6] Laura Wagstaff, Golnar Kolahgar, and Eugenia Piddini. Competitive cell interactions in cancer: a cellular tug of war. Trends in Cell Biology, 23(4):160–167, 2013.

[7] Roghayyeh Baghban, Leila Roshangar, Rana Jahanban-Esfahlan, Khaled Seidi, Abbas Ebrahimi-Kalan, Mehdi Jaymand, Saeed Kolahian, Tahereh Javaheri, and Peyman Zare. Tumor microenvironment complexity and therapeutic implications at a glance. Cell Communication and Signaling, 18(1):59, 2020.

[8] Boris I. Shraiman. Mechanical feedback as a possible regulator of tissue growth. Proceedings of the National Academy of Sciences, 102(9):3318–3323, 2005.

[9] Callie Johnson Miller and Lance A. Davidson. The interplay between cell signalling and mechanics in developmental processes. Nat Rev Genet, 14(10):733–744, 2013.

[10] Henry De Belly, Ewa K. Paluch, and Kevin J. Chalut. Interplay between mechanics and signalling in regulating cell fate. Nat Rev Mol Cell Biol, 23(7):465–480, 2022.

[11] Denis Wirtz, Konstantinos Konstantopoulos, and Peter C. Searson. The physics of cancer: the role of physical interactions and mechanical forces in metastasis. Nat Rev Cancer, 11(7):512–522, 2011.

[12] Edouard Hannezo and Carl-Philipp Heisenberg. Mechanochemical Feedback Loops in Development and Disease. Cell, 178(1):12–25, 2019.

[13] Michael J. Harris, Denis Wirtz, and Pei-Hsun Wu. Dissecting cellular mechanics: Implications for aging, cancer, and immunity. Seminars in Cell & Developmental Biology, 93:16–25, 2019.

[14] M. A. Swartz, D. J. Tschumperlin, R. D. Kamm, and J. M. Drazen. Mechanical stress is communicated between different cell types to elicit matrix remodeling. Proceedings of the National Academy of Sciences, 98(11):6180– 6185, 2001.

[15] Tony Y.-C. Tsai, Mateusz Sikora, Peng Xia, Tugba Colak-Champollion, Holger Knaut, Carl-Philipp Heisenberg, and Sean G. Megason. An adhesion code ensures robust pattern formation during tissue morphogenesis. Science, 370(6512):113–116, 2020.

[16] François Fagotto. Cell sorting at embryonic boundaries. Seminars in Cell & Developmental Biology, 107:126–129, 2020.

[17] Laura Wagstaff, Maja Goschorska, Kasia Kozyrska, Guillaume Duclos, Iwo Kucinski, Anatole Chessel, Lea Hampton-O’Neil, Charles R. Bradshaw, George E. Allen, Emma L. Rawlins, Pascal Silberzan, Rafael E. Carazo Salas, and Eugenia Piddini. Mechanical cell competition kills cells via induction of lethal p53 levels. Nat Commun, 7(1):11373, 2016.

[18] Catarina Brás-Pereira and Eduardo Moreno. Mechanical cell competition. Current Opinion in Cell Biology, 51:15–21, 2018.

[19] Romain Levayer. Solid stress, competition for space and cancer: The opposing roles of mechanical cell competition in tumour initiation and growth. Seminars in Cancer Biology, 63:69–80, 2020.

[20] Jonne Helenius, Carl-Philipp Heisenberg, Hermann E. Gaub, and Daniel J. Muller. Single-cell force spectroscopy. Journal of Cell Science, 121(11):1785–1791, 2008.

[21] Susana Moreno-Flores. Hallmarks of Life in Single Cell Contact Mechanics: Outstanding Challenges and Perspectives. Frontiers in Mechanical Engineering, 6, 2020.

[22] Yuri M. Efremov, Irina M. Zurina, Viktoria S. Presniakova, Nastasia V. Kosheleva, Denis V. Butnaru, Andrey A. Svistunov, Yury A. Rochev, and Peter S. Timashev. Mechanical properties of cell sheets and spheroids: the link between single cells and complex tissues. Biophys Rev, 13(4):541–561, 2021.

[23] M. Krieg, Y. Arboleda-Estudillo, P.-H. Puech, J. Käfer, F. Graner, D. J. Müller, and C.-P. Heisenberg. Tensile forces govern germlayer organization in zebrafish. Nat Cell Biol, 10(4):429–436, 2008.

[24] Wenwei Xu, Roman Mezencev, Byungkyu Kim, Lijuan Wang, John McDonald, and Todd Sulchek. Cell Stiffness Is a Biomarker of the Metastatic Potential of Ovarian Cancer Cells. PLOS ONE, 7(10):e46609, 2012.

[25] Kaoru Sugimura, Pierre-François Lenne, and François Graner. Measuring forces and stresses in situ in living tissues. Development, 143(2):186–196, 2016.

[26] Manuel Gómez-González, Ernest Latorre, Marino Arroyo, and Xavier Trepat. Measuring mechanical stress in living tissues. Nat Rev Phys, 2(6):300–317, 2020.

[27] Vincent Nier, Shreyansh Jain, Chwee Teck Lim, Shuji Ishihara, Benoit Ladoux, and Philippe Marcq. Inference of Internal Stress in a Cell Monolayer. Biophysical Journal, 110(7):1625–1635, 2016.

[28] Ma-lgorzata Lekka, Kajangi Gnanachandran, Andrzej Kubiak, Tomasz Zieliński, and Joanna Zemla. Traction force microscopy – Measuring the forces exerted by cells. Micron, 150:103138, 2021.

[29] Thuan Beng Saw, Amin Doostmohammadi, Vincent Nier, Leyla Kocgozlu, Sumesh Thampi, Yusuke Toyama, Philippe Marcq, Chwee Teck Lim, Julia M. Yeomans, and Benoit Ladoux. Topological defects in epithelia govern cell death and extrusion. Nature, 544(7649):212–216, 2017.

[30] Vincent Nier, Grégoire Peyret, Joseph d’Alessandro, Shuji Ishihara, Benoit Ladoux, and Philippe Marcq. Kalman Inversion Stress Microscopy. Biophysical Journal, 115(9):1808–1816, 2018.

[31] Grégoire Peyret, Romain Mueller, Joseph d’Alessandro, Simon Begnaud, Philippe Marcq, Reńe-Marc Mège, Julia M. Yeomans, Amin Doostmohammadi, and Benoît Ladoux. Sustained Oscillations of Epithelial Cell Sheets. Biophysical Journal, 117(3):464–478, 2019.

[32] Surabhi Sonam, Lakshmi Balasubramaniam, Shao-Zhen Lin, Ying Ming Yow Ivan, Irina Pi-Jaumà, Cecile Jebane, Marc Karnat, Yusuke Toyama, Philippe Marcq, Jacques Prost, Reńe-Marc Mège, Jean-François Rupprecht, and Benoît Ladoux. Mechanical stress driven by rigidity sensing governs epithelial stability. Nat. Phys., 19(1):132–141, 2023.

[33] Keisuke Kuromiya, Kana Aoki, Kojiro Ishibashi, Moe Yotabun, Miho Sekai, Nobuyuki Tanimura, Sayuri Iijima, Susumu Ishikawa, Tomoko Kamasaki, Yuki Akieda, Tohru Ishitani, Takashi Hayashi, Satoshi Toda, Koji Yokoyama, Chol Gyu Lee, Ippei Usami, Haruki Inoue, Ichigaku Takigawa, Estelle Gauquelin, Kaoru Sugimura, Naoya Hino, and Yasuyuki Fujita. Calcium sparks enhance the tissue fluidity within epithelial layers and promote apical extrusion of transformed cells. Cell Reports, 40(2):111078, 2022.

[34] Eduardo Moreno, Léo Valon, Florence Levillayer, and Romain Levayer. Competition for Space Induces Cell Elimination through Compaction-Driven ERK Downregulation. Current Biology, 29(1):23–34.e8, 2019.

[35] Eduardo Moreno, Konrad Basler, and Gińes Morata. Cells compete for Decapentaplegic survival factor to prevent apoptosis in Drosophila wing development. Nature, 416(6882):755–759, 2002.

[36] Masakazu Hashimoto and Hiroshi Sasaki. Cell competition controls differentiation in mouse embryos and stem cells. Current Opinion in Cell Biology, 67:1–8, 2020.

[37] Mihoko Kajita, Kaoru Sugimura, Atsuko Ohoka, Jemima Burden, Hitomi Suganuma, Masaya Ikegawa, Takashi Shimada, Tetsuya Kitamura, Masanobu Shindoh, Susumu Ishikawa, Sayaka Yamamoto, Sayaka Saitoh, Yuta Yako, Ryosuke Takahashi, Takaharu Okajima, Junichi Kikuta, Yumiko Maijima, Masaru Ishii, Masazumi Tada, and Yasuyuki Fujita. Filamin acts as a key regulator in epithelial defence against transformed cells. Nat Commun, 5(1):4428, 2014.

[38] Joseph D. Dukes, Paul Whitley, and Andrew D. Chalmers. The MDCK variety pack: choosing the right strain. BMC Cell Biology, 12(1):43, 2011.

[39] Manuel Théry and Matthieu Piel. Micropatterning in Cell Biology, Part C. Number pt. 3 in Methods in Cell Biology. Elsevier Science, 2014.

[40] Thomas E. Angelini, Edouard Hannezo, Xavier Trepat, Manuel Marquez, Jeffrey J. Fredberg, and David A. Weitz. Glass-like dynamics of collective cell migration. Proceedings of the National Academy of Sciences, 108(12):4714–4719, 2011.

[41] M S Steinberg and M Takeichi. Experimental specification of cell sorting, tissue spreading, and specific spatial patterning by quantitative differences in cadherin expression. Proceedings of the National Academy of Sciences, 91(1):206–209, 1994.

[42] Tetsuhisa Otani, Thanh Phuong Nguyen, Shinsaku Tokuda, Kei Sugihara, Taichi Sugawara, Kyoko Furuse, Takashi Miura, Klaus Ebnet, and Mikio Furuse. Claudins and JAM-A coordinately regulate tight junction formation and epithelial polarity. Journal of Cell Biology, 218(10):3372–3396, 2019.

[43] Lakshmi Balasubramaniam, Amin Doostmohammadi, Thuan Beng Saw, Gautham Hari Narayana Sankara Narayana, Romain Mueller, Tien Dang, Minnah Thomas, Shafali Gupta, Surabhi Sonam, Alpha S. Yap, Yusuke Toyama, Reńe-Marc Mège, Julia M. Yeomans, and Benoît Ladoux. Investigating the nature of active forces in tissues reveals how contractile cells can form extensile monolayers. Nat. Mater., 20(8):1156–1166, 2021.

[44] M. C. Marchetti, J. F. Joanny, S. Ramaswamy, T. B. Liverpool, J. Prost, Madan Rao, and R. Aditi Simha. Hydro-dynamics of soft active matter. Rev. Mod. Phys., 85:1143–1189, 2013.

[45] De-Qing Zhang, Peng-Cheng Chen, Zhong-Yi Li, Rui Zhang, and Bo Li. Topological defect-mediated morphodynamics of active–active interfaces. Proceedings of the National Academy of Sciences, 119(50):e2122494119, 2022.

[46] Pengtao Yue, James J. Feng, Chun Liu, and Jie Shen. A diffuse-interface method for simulating two-phase flows of complex fluids. Journal of Fluid Mechanics, 515:293–317, 2004.

[47] Fernando Caballero and M. Cristina Marchetti. Activity-suppressed phase separation. Phys. Rev. Lett., 129:268002, 2022.

[48] Katharina P. Landsberg, Reza Farhadifar, Jonas Ranft, Daiki Umetsu, Thomas J. Widmann, Thomas Bittig, Amani Said, Frank Jülicher, and Christian Dahmann. Increased Cell Bond Tension Governs Cell Sorting at the Drosophila Anteroposterior Compartment Boundary. Current Biology, 19(22):1950–1955, 2009.

[49] Floris Bosveld, Boris Guirao, Zhimin Wang, Mathieu Rivière, Isabelle Bonnet, François Graner, and Yohanns Bellaïche. Modulation of junction tension by tumor suppressors and proto-oncogenes regulates cell-cell contacts. Development, 143(4):623–634, 2016.

[50] Praver Gupta, Sayantani Kayal, Shilpa P. Pothapragada, Harish K. Senapati, Padmashree Devendran, Dapeng Bi, and Tamal Das. Mechanical imbalance between normal and cancer cells drives epithelial defense against cancer. bioRxiv, 2023.

[51] Oren Shaya, Udi Binshtok, Micha Hersch, Dmitri Rivkin, Sheila Weinreb, Liat Amir-Zilberstein, Bassma Khamaisi, Olya Oppenheim, Ravi A. Desai, Richard J. Goodyear, Guy P. Richardson, Christopher S. Chen, and David Sprinzak. Cell-Cell Contact Area Affects Notch Signaling and Notch-Dependent Patterning. Developmental Cell, 40(5):505–511.e6, 2017.

[52] Léo Guignard, Ulla-Maj Fiúza, Bruno Leggio, Julien Laussu, Emmanuel Faure, Gaël Michelin, Kilian Biasuz, Lars Hufnagel, Grégoire Malandain, Christophe Godin, and Patrick Lemaire. Contact area–dependent cell communication and the morphological invariance of ascidian embryogenesis. Science, 369(6500):eaar5663, 2020.

[53] Laura Canty, Eleyine Zarour, Leily Kashkooli, Paul François, and François Fagotto. Sorting at embryonic boundaries requires high heterotypic interfacial tension. Nat Commun, 8(1):157, 2017.

[54] Tetsuo Ichii and Masatoshi Takeichi. p120-catenin regulates microtubule dynamics and cell migration in a cadherin-independent manner. Genes to Cells, 12(7):827–839, 2007.

[55] Yasuto Takeuchi, Rika Narumi, Ryutaro Akiyama, Elisa Vitiello, Takanobu Shirai, Nobuyuki Tanimura, Keisuke Kuromiya, Susumu Ishikawa, Mihoko Kajita, Masazumi Tada, Yukinari Haraoka, Yuki Akieda, Tohru Ishitani, Yoichiro Fujioka, Yusuke Ohba, Sohei Yamada, Yoichiroh Hosokawa, Yusuke Toyama, Takaaki Matsui, and Yasuyuki Fujita. Calcium Wave Promotes Cell Extrusion. Current Biology, 30(4):670–681.e6, 2020.

[56] Shunsuke Kon, Kojiro Ishibashi, Hiroto Katoh, Sho Kitamoto, Takanobu Shirai, Shinya Tanaka, Mihoko Kajita, Susumu Ishikawa, Hajime Yamauchi, Yuta Yako, Tomoko Kamasaki, Tomohiro Matsumoto, Hirotaka Watanabe, Riku Egami, Ayana Sasaki, Atsuko Nishikawa, Ikumi Kameda, Takeshi Maruyama, Rika Narumi, Tomoko Morita, Yoshiteru Sasaki, Ryosuke Enoki, Sato Honma, Hiromi Imamura, Masanobu Oshima, Tomoyoshi Soga, Jun-ichi Miyazaki, Michael R. Duchen, Jin-Min Nam, Yasuhito Onodera, Shingo Yoshioka, Junichi Kikuta, Masaru Ishii, Masamichi Imajo, Eisuke Nishida, Yoichiro Fujioka, Yusuke Ohba, Toshiro Sato, and Yasuyuki Fujita. Cell competition with normal epithelial cells promotes apical extrusion of transformed cells through metabolic changes. Nat Cell Biol, 19(5):530–541, 2017.

[57] Kenry, Man Chun Leong, Mui Hoon Nai, Fook Chiong Cheong, and Chwee Teck Lim. Viscoelastic Effects of Silicone Gels at the Micro- and Nanoscale. Procedia IUTAM, 12:20–30, 2015.

[58] Maimon M. Cohen and Margery W. Shaw. Effects of Mitomycin C on Human Chromosomes. Journal of Cell Biology, 23(2):386– 395, 1964.

[59] Stephan Preibisch, Stephan Saalfeld, and Pavel Tomancak. Globally optimal stitching of tiled 3D microscopic image acquisitions. Bioinformatics, 25(11):1463–1465, 2009.

[60] Jean-Louis Martiel, Aldo Leal, Laetitia Kurzawa, Martial Balland, Irene Wang, Timothée Vignaud, Qingzong Tseng, and Manuel Théry. Chapter 15 - Measurement of cell traction forces with ImageJ. In Ewa K. Paluch, editor, Methods in Cell Biology, volume 125, pages 269–287. Academic Press, 2015.

[61] F. Hecht. New development in freefem++. J. Numer. Math., 20(3-4):251–265, 2012.

